# Development of BODIPY-based fluorescent probes for highly selective amino acid identification

**DOI:** 10.1101/2024.04.02.587799

**Authors:** Nathaniel Hendrick, Andrew Martin, Andrew Chan, Jason Lenihan, Harrison Reiter, Mitchell Clough, Lian Jiang, Eddie Whaley, Yoo Jeong Huh, James McNeely, Anderson Chen, Andrew Emili, Aaron Beeler

## Abstract

Amino acid sensing is a powerful tool for intra- and extra-cellular assays. We report herein the generation and optimization of bioconjugatable fluorescent reporter for pan-amino acid sensing, the OMNI probe, that through inductive electronics produces a unique photophysical response upon coupling to the α-amino group of an amino acid depending on the identity of the adjacent side chain. Using an auto-photocatalyzed Meerwein arylation reaction, we created a library of bis-arylated 8-methylthio-BODIPY dye derivatives exhibiting improved spectral variation, brightness, dynamic range, robustness, water solubility, and reactivity. Aryl groups with optimal photophysical behaviors were combined to create asymmetric versions conferring high spectral range, brightness, and water solubility. Combining aryl group modifications yielded a set of engineered asymmetric probes that achieved complete spectral resolution in a multi-variable analysis (fluorescent emission, lifetime, brightness) for all twenty native amino acid-dye conjugates while increasing reactivity toward amino acids in aqueous buffer. With our robust OMNI probe design, pan-amino acid resolution and identification through high sensitivity fluorescence down to single molecules is now routinely possible for diverse potential applications including protein sequencing.

## INTRODUCTION

Amino acids are key biomolecules for cellular function, and are essential for protein synthesis, cell signaling, and metabolic regulation in healthy cells.^1-3^ Dysregulation of amino acid production, usage or accumulation within a cell can impact a myriad of biochemical processes, and is associated with diverse diseased states.^3-5^ Thus, developing tools and assays to detect and classify the presence each of the twenty native amino acids in a sample is of great importance to bioanalytical chemistry, as it can help with phenotyping cellular states.^6^

Primary methods for identifying amino acids involve mass spectrometry (MS), high performance liquid chromatography (HPLC), or capillary electrophoresis (CE). Amino acids are typically identified using these methods by matching their characteristic retention times or mass spectral features .^5,7-9^ The advantages of these classical methods are the specific identification of certain amino acids, and ability to quantify their respective concentrations within a given sample. However, the modest sensitivity and detection limits of existing methods challenge their ability to accurately detect single molecule concentrations.

Innovations in solid-state and biological nanopores have enabled the identification of most, if not all, twenty naturally occurring amino acids with high confidence, even at single micromolar concentrations.^10,11^ Despite their effectiveness, the specialized equipment and expertise required for nanopore fabrication limit their accessibility to many research groups. We envisioned developing a universal fluorescent probe suitable for readout on standard laboratory equipment, that can unambiguously identify and quantify all naturally occurring amino acids through unique spectral responses.

Fluorescent probes have been previously designed for the identification of specific amino acids via bioconjugation, facilitating targeted intra- and extracellular detection. These strategies have enabled the detection of single molecules within the complex spatial environment of cells.^11-17^ The current range of probes is limited, typically identifying only one or two of the twenty native amino acids, with a preference for coupling to specific side chain residues. This limitation restricts the breadth of information obtainable from any single probe.

Farinone et al. highlighted the potential of 8-methylthio- BODIPY (Biellmann BODIPY) to react with α-amines and nucleophilic side chains of amino acids under mild conditions, albeit with low efficiency.^18^ They noted distinct spectral responses post-conjugation, suggesting the dye’s sensitivity to minor structural variations in the bound amine. Building on this premise, we explored whether 8- methylthio-BODIPY and its derivatives could be tailored to react more efficiently with amino acid N-termini, enabling differentiation between various amino acid side chains in aqueous buffers compatible with biological systems..

## RESULTS

### Dye Synthesis

We investigated the ability of 8-methylthio-BODIPY to provide spectral resolution for all twenty naturally occurring amino acids, utilizing fluorescent emission as the readout. The compound, 8-methylthio-BODIPY (**1**) (**Figure 1a**) was synthesized according to literature procedures and reacted with the amino acids in solution at neutral pH.^19^ Remarkably, we observed that each amino acid-dye conjugate (AADC) exhibited distinct emission spectra, characterized by unique profile shapes and peak maxima. These conjugates demonstrated fully resolvable Stokes shifts across a dynamic range of 13 nm (Figure 1b). This prompted a deeper exploration into this particularly responsive class of fluorescent molecules, which could serve as a promising scaffold for amino acid detection.

**Figure 1.**
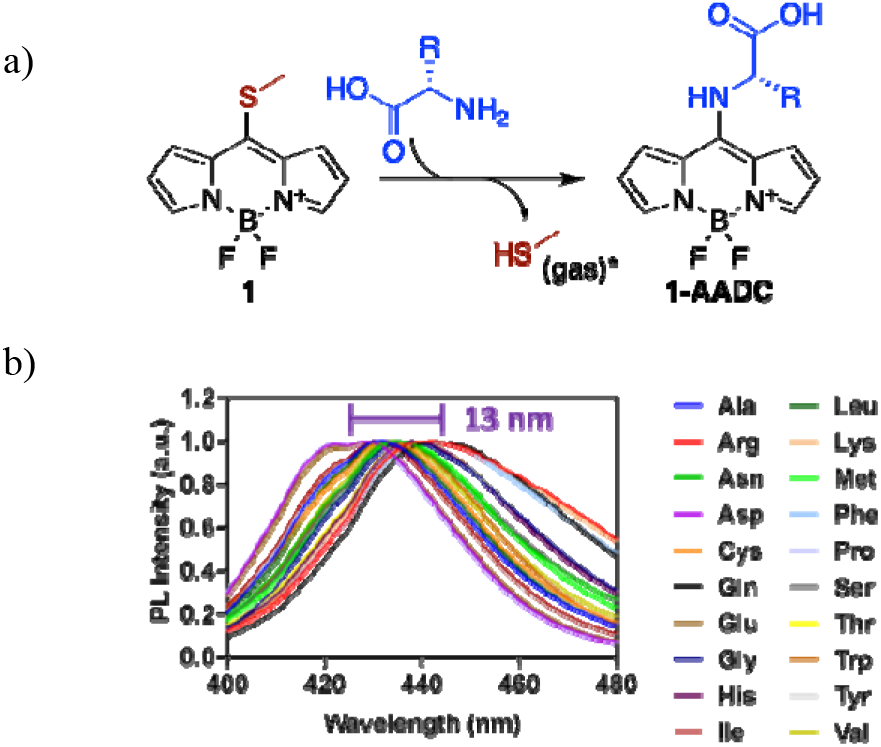
a) Scheme of **1** reacting with an amino acid. b) Normalized emission spectra of AADCs reacted with 1. All measurements were taken in 1 M Tris buffer.

First, we sought to understand the structural properties of the fluorophore that conferred a sizeable dynamic range between AADCs. Moreover, we aimed to identify other biophysical factors influencing probe reactivity, such as water solubility, selective coupling to amines, and fluorescence intensity in aqueous environments. Based on the commercial availability of substituted aryl-starting materials, we envisioned that arylation of the BODIPY α-positions would allow us create a biaryl dye would provide a versatile empirical method for exploring which properties lead to a desirable sensor.

Our initial attempt at synthesis of biaryl dyes started with formation of a biaryl pyrrole using pyrrole-boronic acid and bromobenzene via a Suzuki coupling.^20^ This step gave modest yields and the resulting thioketone formation and boron complexation were less efficient compared to pyrrole alone, a situation we anticipated would worsen with addition of electron withdrawing groups on the phenyl rings. We also reasoned that diversifying at such an early stage would greatly increase the number of synthetic steps and purifications needed to build a small library of dyes. Consequently, we searched for alternate late-stage arylation techniques beginning with a common starting materials. Previously, Peña-Cabrera et al had reported late-stage arylation of 8-methylthio-BODIPYs using in situ Meerwein arylation, but obtained a single bis-arylated molecule in low yield.^21^ In contrast, Jiao et al achieved higher yields of late-stage bis-arylation of 8-aryl-BODIPY molecules using an auto-photocatalyzed Meerwein arylation with aryl diazonium salts.^22^ By employing the auto-photocatalyzed Meerwein arylation reaction, we efficiently synthesized a collection of symmetrical bis-arylated BODIPY dyes. These dyes feature various electron- donating and withdrawing groups positioned around the phenyl rings (**Figure 2**), all derived from a single common precursor (**1**).

**Figure 2.**
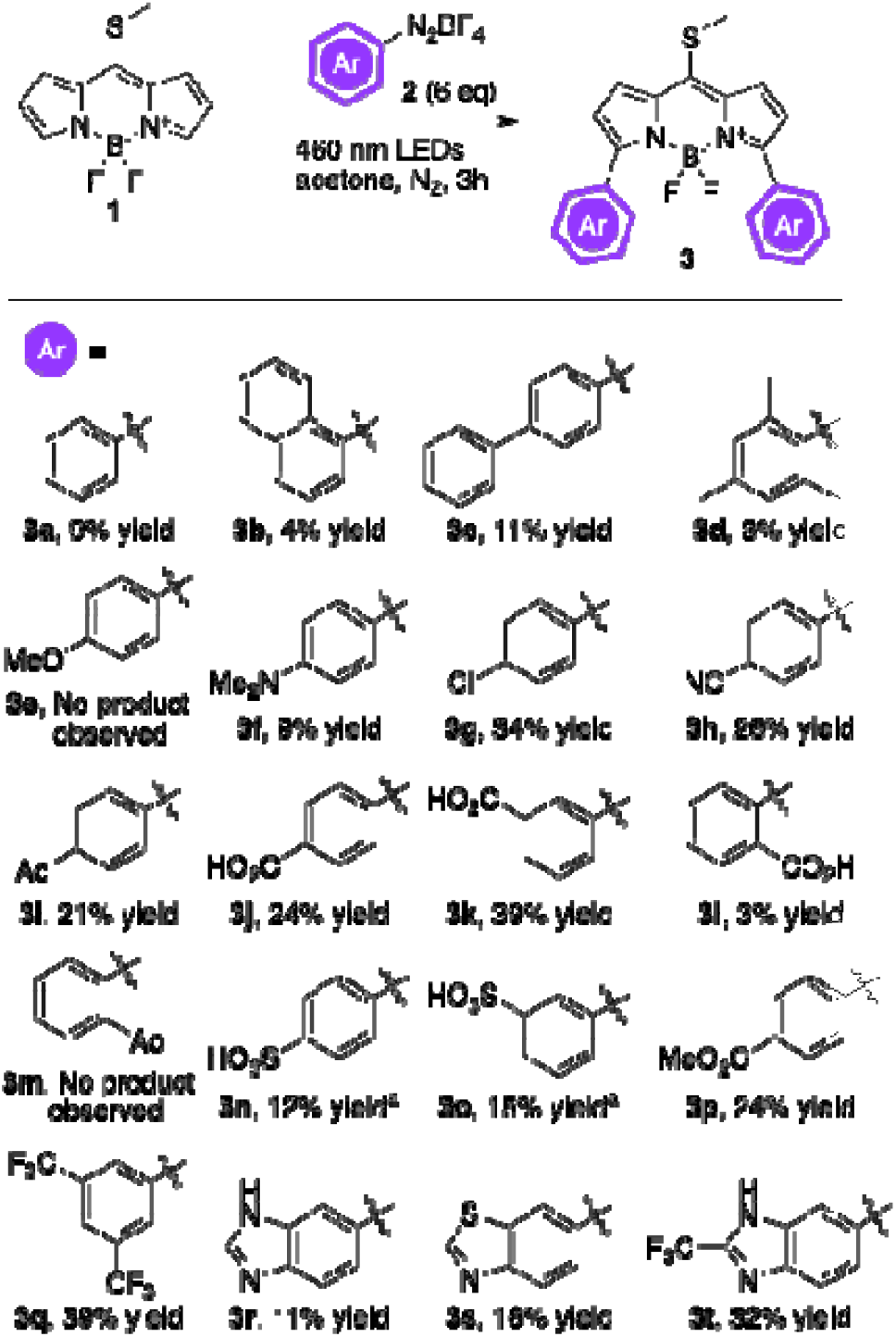
Scheme of auto-photocatalyzed Meerwein arylation. Reactions were conducted at ambient temperature at 100 mM. ^a^Reaction was conducted in 25% DMSO/acetone.

Compounds with electron donating groups displayed lower yields whereas electron withdrawing groups at the para- or meta-positions showed the highest yields (**Figure 2**), likely due to stabilization of the aryl radical intermediates. Attempts to add aryl groups via ortho-substitution resulted in the lowest conversion, likely due to steric effects as the drop in yield occurred with both electron donating and withdrawing groups.

### Amino Acid-Dye Conjugate Testing

With a starting set of bis-arylated dyes in hand, we tested their reactivity with amino acids and analyzing the conjugated products to assess their photophysical responses. Each dye was coupled with all twenty naturally occurring amino acids in a pH 9.2 bicarbonate buffer for 16 h, and then diluted in 1 M Tris buffer to a final concentration of 1 μM to quench unreacted dye. Surprisingly, some of the dyes tested showed limited reaction when incubated with amino acids, likely due to limited solubility in aqueous media. Dyes **3b, 3c, 3d, 3g**, and **3q**, visibly precipitated out of solution, and had little observable reaction. The ortho- substituted 3d and 3l were fully soluble but, steric encumbrance of the ortho-substitution did not not allow reaction of the amine. The most reactive constructs had electron withdrawing groups at the para-positions of the aryl substitutions such as **3h-j, 3n**, and **3p**. while the presence of electron donating groups slowed down dye reactivity. As expected, inductive effects of meta-substituted electron withdrawing groups showed little effect on reaction time. Notably, sulfonate substituted **3n** and **3o** displayed excellent water solubility and readily dissolved in water without precipitation.

We systematically assessed the spectral response and brightness of each AADC in aqueous buffer (**Figure 3a**). From these measurements, we elucidated a structure- activity relationship for the dye constructs. Notably, while almost all bis-arylated AADCs exhibited a spectral range greater than dye **1** (**Figure 3b**), confirming our initial rationale for arylation, their photophysical response did not show a consistent trend between different dye congeners with the same amino acid; one amino acid could be the most red-shifted for one dye but not for another. This unexpected result is likely due to a combination of the intrinsic sensitivity of the BODIPY core structure to subtle inductive electronic changes and through-space interactions between the amino acid side chain and the aryl substitution.

**Figure 3.**
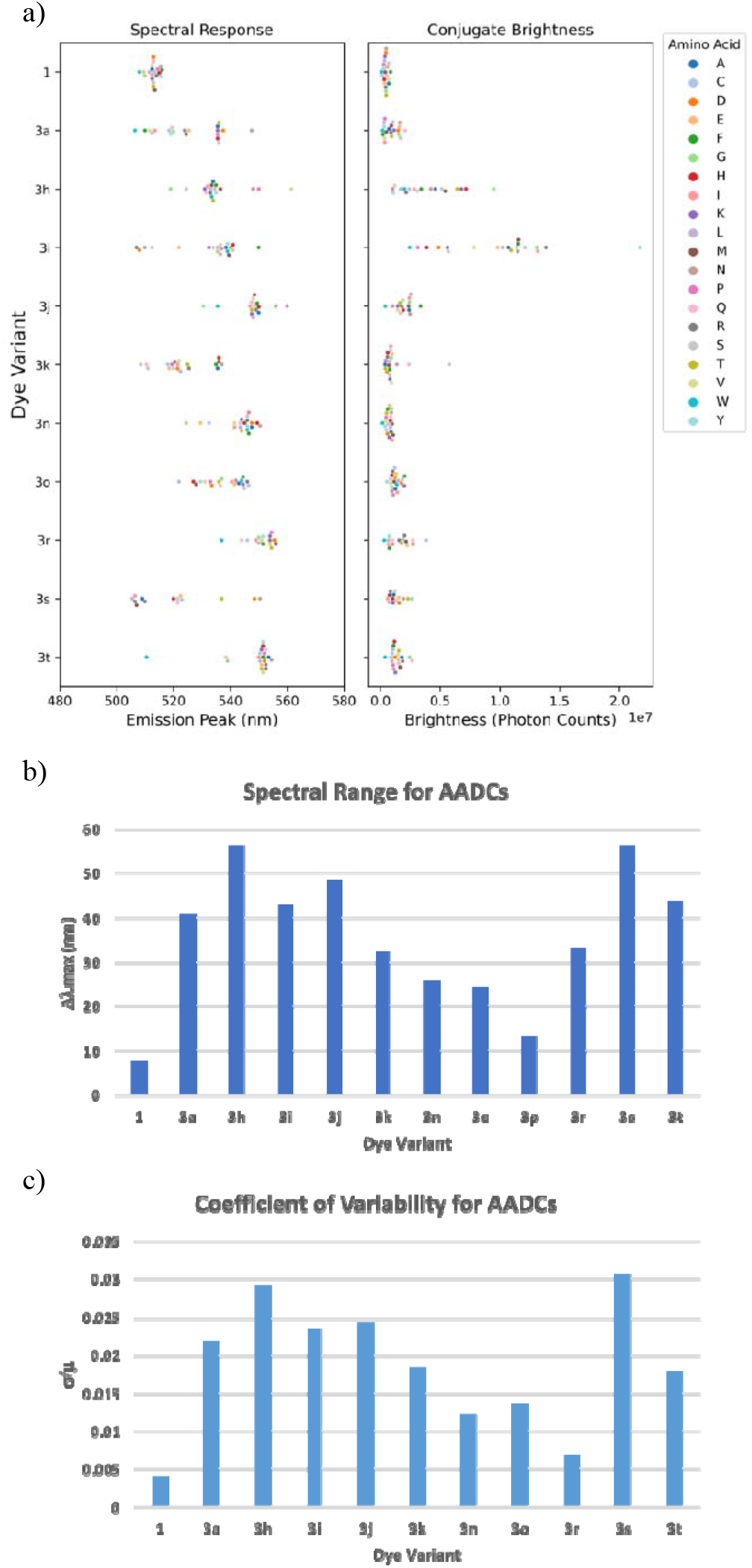
a) Spectral analyses of AADCs made with bisarylated dyes 3. All measurements taken with 1 □M solutions in 1 M Tris buffer excited at 440 nm. (*Left*) Maximal emission recorded for each 3-AADC. (*Right*) Photon count for each 3- AADC. b) Spectral range determined based on difference between reddest to bluest emitting AADC within a single dye construct. c) Coefficient of variation in emission peak for all 20 native amino acids within each 3-AADC construct.

Examination of the distribution of the AADC spectral response revealed that certain dyes displayed a large dynamic range (lowest to highest emission maxima), but the majority clustered with similar emission patterns, such as **3n, 3r**, and **3t** especially. The latter result was undesirable because while a handful of amino acids could easily be identified, a more even distribution between AADCs is warranted to achieve pan-amino acid resolution for more confident identification. We determined the coefficient of variation for each set of AADCs to quantify the distribution for each dye (**Figure 3c**). From this coefficient, we were able to gauge and compare the overall distributions of AADCs between the dye constructs. Notably, benzothiazole (**3s**) and *p-*cyano (**3h**) biaryl derivatives displayed the greatest variation between AADCs, but other electron withdrawing groups including acyl (**3i**) or carboxylic (**3j**) groups also conferred high coefficients of variation. In addition, dyes **3i** and **3j** displayed the highest conjugate brightness in aqueous media out of all symmetrical dyes tested. While these dyes displayed excellent spectral properties, they were still insoluble in aqueous buffer which decreased their ability to efficiently react with naturally occurring amino acids without an organic cosolvent.

### Non-symmetrical Dye Synthesis

Given the limitations of most symmetrical derivatives, we sought to leverage the bis-arylated BODIPY properties further by combining higher-performing aryl groups. By making asymmetric bis-arylated BODIPY dyes with a water-solubilizing arm and a spectral separation arm, we investigated if these properties were combinatorial. While synthesis of the asymmetric dyes proceeded through a similar route as for their symmetrical counterparts, we reduced the aryl diazonium salt equivalents by half to obtain mono-arylation products. The mono-arylated products were then reacted again with a different aryl diazonium salt to create a final asymmetric bis-arylated dye (**Figure 4a**). As the water-solubilizing arm tested *m*- and *p*- sulfonates and *p-*carboxylic acid, as all three showed high solubility in symmetrical constructs, we crossed those arms with *p*-acyl, *p-*cyano, and benzothiazole that gave the highest spectral discrimination between AADCs (resolution) and highest overall photon counts (brightness) (**Figure 4b**).

**Figure 4.**
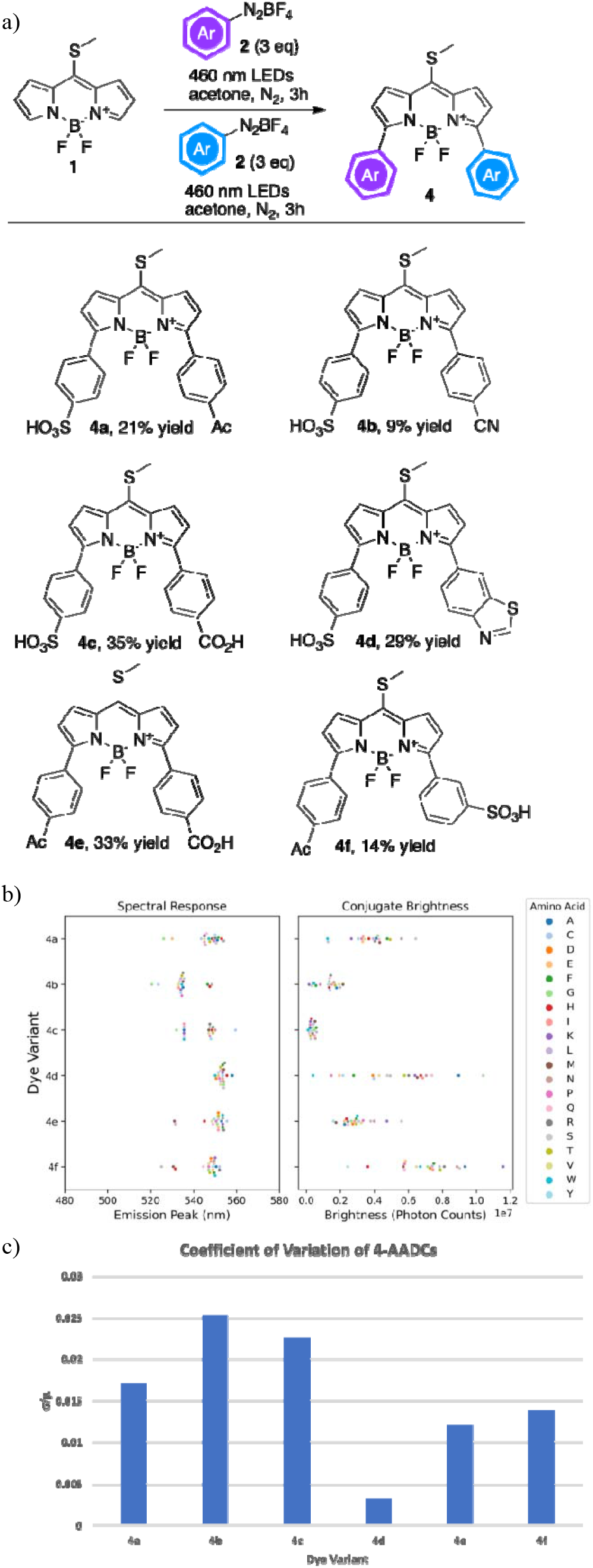
a) Scheme of auto-photocatalyzed Meerwein arylation. b) Spectral analyses of AADCs made with bisarylated dyes **4**. All measurements taken as 1 μM solutions (in 1 M Tris buffer) after excitation at 440 nm. (*Left*) Emission max and (*Right*) photon count measured for each 4- AADC. b) Coefficient of variation for all amino acid recorded within each 4-AADC construct.

As expected, all asymmetrical dyes tested were highly soluble in water, and highly reactive to the amino acids when conjugated using the same conditions used for the symmetrical AADC testing. Moreover, when comparing the spectral range and coefficient of variation for the asymmetric dyes (**Figure 4c**), the values for the dual electron withdrawing arms **4a-c** and **4e-f** were the same or only slightly less than their symmetrical versions. In contrast, **4d** completely lost spectral range and variation, likely due to the push-pull nature of electrons from the electron donating benzothiazole arm to the electron withdrawing sulfonate arm reducing the influence of the amino acid side chain on the overall conjugated system.

We found that the *p-*acyl dyes **4a** and **4f** displayed the best overall spectral range, AADC distribution, and brightness in aqueous buffer. Coupled with their good water solubility, these dyes showed the greatest ability to pan-identify amino acids under mild, aqueous conditions. Notably, including other spectral information such as fluorescent lifetime and photon counts as second and third dimensions for identification allowed for greater resolution between amino acids that produced spectrally similar emission profiles. For example, when comparing the fluorescent lifetime of **4a-AADCs** versus peak emission, a high degree of separation between individual amino acids was achieved (**Figure 5a**), which could be even further enhanced by factoring in relative brightness (**Figure 5b**).

**Figure 5.**
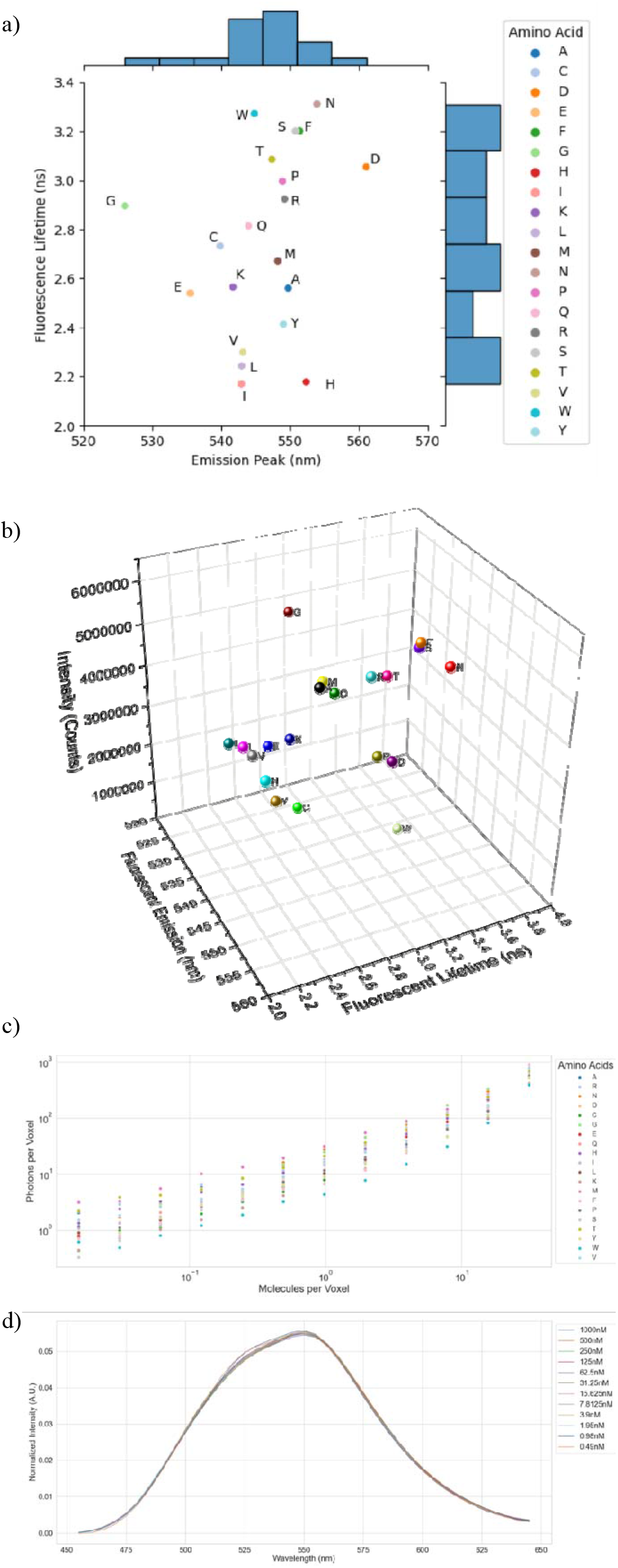
a) Normalized two-dimensional plot with corresponding histograms of fluorescent emission (nm) versus fluorescent lifetime (ns) for **4a** reacted with all twenty amino acids. All measurements taken at 1 μM in 1 M Tris buffer. b) Normalized 3D plot of fluorescent emission (nm) versus fluorescent lifetime (ns) and fluorescent intensity (photon counts) for 4a reacted with all twenty amino acids (1 μM in 1 M Tris buffer. c) Plot of photons per voxel versus molecules per voxel of **4a** reacted with all twenty amino acids at varying concentrations (in 1 M Tris buffer). d) Normalized emission spectra of 4a reacted with alanine at varying concentrations (in 1 M Tris buffer).

Hence, using routine multi-modal fluorescent readouts, we obtained probes that could confidently identify amino acids in solution based on their discrete photophysical properties. A serial dilution study was performed using **4a- AADCs** to measure signal and spectral response at lower concentrations. All AADCs of **4a** showed good intensity down to single molecule (or lower) per voxel concentrations (**Figure 5c**). In addition, spectral shape and emission max of the AADCs was maintained as well with alanine (**Figure 5d**).

## DISCUSSION

Synthesis of bis-arylated 8-methylthio-BODIPY dye derivatives proved fruitful basis for generating probes that can spectrally resolve all twenty natural amino acids. Through an auto-photocatalyzed Meerwein reaction, we created a library of bis-arylated amino reactive dyes and explored the structure-activity relationship of amino acid-dye conjugates. From this exploration, we combined desirable features into a novel set of asymmetric bis-arylated dyes, which consist of two different aryl arms conferring complementary attributes. This combinatorial strategy allowed us to define the biophysically unique probes that conferred sufficient spectral resolution and variation between AADCs, while also increasing water solubility and brightness.

Our optimized OMNI probe (**4a**) displayed good spectral intensity with amino acid conjugates down to sub- nanomolar concentrations, and maintained distinct and identifiable spectral responses for different amino acids. Thus, OMNI probe can unambiguously identify individual amino acids at single-molecule resolution. In terms of potential applications, we are actively exploring the potential utility of using these probes to sequentially deduce the N- terminal residues and hence sequences of cellular proteins with the same confidence as detection of single amino acids.

## METHODS

### Dye synthesis and characterization

Preparation of compound **1** was synthesized according to literature procedure.^19^ Experimental protocols for the synthesis of bis-arylated 8-methylthio-BODIPY dyes and their corresponding characterization data are provided in the *Supplementary Information*.

### Amino acid-dye conjugation

Prior to conjugation, all dyes were made into 10 mM stock solutions in DMSO. In a 1536 polypropylene well plate, 1 μL of 10 mM dye stock solution was added to 6 μL 100 mM pH 9.2 sodium bicarbonate buffer followed by 3 μL 50 mM amino acid solution in deionized water. This dilution resulted in the final concentration of 1 mM of dye in each well. Twenty wells were made for each dye tested containing each naturally occurring amino acid. After addition of all reagents to their respective wells, the multi-well plate was sealed with PCR film and stirred on a BioShake iQ plate shaker at 2000 rpm at ambient temperature for 16 hours. After 16 hours, most dyes were fully consumed and conjugated to the amino acids. There is a stark optical shift of the solution from pink to yellow-green upon full reaction of the dye with the amino acids. 1 μL of each of the reacted AADCs were pipetted into 99 μL of 1 M pH 7.6 Tris buffer in a 384-glass bottom well plate. This dilution changes the concentration of the AADC to 10 μM. 95 μL of each AADC solution was discarded, and 45 μL of 1 M pH 7.6 Tris buffer was added back to each well making the final concentration of AADC at 1 μM.

### Spectral and lifetime imaging in solution

Samples containing the dye conjugated with monomers were pipetted into 384 well glass bottom plates with a #1.5 cover glass. The imaging of these plates was performed with a Leica SP8 confocal microscope with a white light laser (WLL) source, 20x 0.75 NA objective (Model: HC PL APO CS2), and the FALCON time correlated single photon counting (TCSPC) fluorescence lifetime addon. For all the BODIPY monomer conjugates, 440nm laser excitation, at 85.4μW with a repetition rate of 80MHz was used to generate fluorescence photons from the dye. A 445nm, 10nm bandwidth notch filter is used to suppress contamination of the fluorescence spectra by reflection from the laser excitation source. The fluorescence spectrums of the dye conjugates were acquired over a 1502μm^2^ region of interest, at 128px by 128px, 30.78μs dwell time, at 10nm optical bandwidth and 5nm step sizes over the 450nm to 650nm range. To remove the Raman signal from our solution measurements, spectra from wells containing buffer solution only were also acquired and then subtracted from the measured dye-conjugate spectra. For fluorescence lifetime measurements, the arrival times of the photons were counted using 450nm-650nm fluorescence spectral range with the same laser excitation wavelength as the spectral scan, but at 20MHz repetition rate and 9.41μW. The lower power eliminates photon pileup errors and the lower repetition rate better captures the full-time binned fluorescence decay curve. The resulting time binned fluorescence decay data was fitted to a tri-exponential decay curve, and the mean intensity-weighted lifetime was extracted for each sample. All measurements were performed with the same hybrid detector on the microscope.

### Extraction and analysis of solution data

Following the image acquisition, a .lif file is generated from the LAS X software which contains the raw data for the images. Within each 384 well plate, a subsection of the wells was allocated to contain the buffer solution used to dilute the other samples within the plate. These wells were used to calculate any background signals that are present when imaging the solution, such as Gaussian noise and Raman peaks. The average of these buffer wells was calculated, and this average was then subtracted pixel-wise from all of the sample wells within the same plate. This pixel-wise subtraction allowed the removal of background noise from the solution-based samples. Following this denoising step, 5-by-5 super-pixels were generated for each image to improve the signal-to-noise ratio. To further remove Gaussian noise, the spectra for each image was smoothed using a Savitzky-Golay filter. After some initial testing, a window length of 9 and a polynomial to the 3rd order was found to be the most optimal. For each sample well, 5 regions of interest (ROIs) were imaged, and thus calculated the average smoothed spectra for each sample from these ROIs. Subsampling was also performed on each sample’s spectra using 1nm intervals to more accurately determine the peak wavelength of a sample. Finally, the peak wavelength is extracted from the merged, smoothed, and subsampled spectra of each sample. In addition, the total photon counts for each sample is extracted by summing all of the pixel values within the images.

## ASSOCIATED CONTENT

### Supporting Information

Synthesis and characterization data, photophysical data, theoretical calculations

## AUTHOR INFORMATION

### Notes

Any additional relevant notes should be placed here.

## ACKNOWLEDGMENT

(Word Style “TD_Acknowledgments”). Generally the last paragraph of the paper is the place to acknowledge people (dedications), places, and financing (you may state grant numbers and sponsors here). Follow the journal’s guidelines on what to include in the Acknowledgement section.

